# Physical mechanisms of ESCRT-III-driven cell division in archaea

**DOI:** 10.1101/2021.03.23.436559

**Authors:** L. Harker-Kirschneck, A. E. Hafner, T. Yao, A. Pulschen, F. Hurtig, C. Vanhille-Campos, D. Hryniuk, S. Culley, R. Henriques, B. Baum, A. Šarić

**Affiliations:** Department of Physics & Astronomy, UCL, London, United Kingdom; Institute for the Physics of Living Systems, UCL, London, United Kingdom; MRC Laboratory for Molecular Cell Biology, UCL, London, United Kingdom; MRC Laboratory Molecular Biology, Cambridge, United Kingdom

## Abstract

Living systems propagate by undergoing rounds of cell growth and division. Cell division is at heart a physical process that requires mechanical forces, usually exerted by protein assemblies. Here we developed the first physical model for the division of archaeal cells, which despite their structural simplicity share machinery and evolutionary origins with eukaryotes. We show how active geometry changes of elastic ESCRT-III filaments, coupled to filament disassembly, are sufficient to efficiently split the cell. We explore how the non-equilibrium processes that govern the filament behaviour impact the resulting cell division. We show how a quantitative comparison between our simulations and dynamic data for ESCRTIII-mediated division in *Sulfolobus acidocaldarius*, the closest archaeal relative to eukaryotic cells that can currently be cultured in the lab, and reveal the most likely physical mechanism behind its division.

---

Cell division is one of the most fundamental requirements for the existence of life on Earth. During division the material from a single cell is divided into two separate daughter cells. This is an inherently physical process. Living cells have evolved multiple ways to apply mechanical forces for this purpose. In general, the process involves proteins that assemble into long polymeric filaments at the cytoplasmic side of the cell membrane. The filaments then undergo a series of energy-driven changes in their form and organisation to deform the associated membrane and/or guide cell wall assembly. The physics of such non-equilibrium protein self-assembly processes that produce the mechanical work needed to reshape and cut soft surfaces is largely underexplored.

Although the mechanisms of division differ across the tree of life, recent data support the idea that eukaryotic cells likely arose from the symbiosis of an archaeal cell and an alphaproteobacterial cell, where the archaeal host gave rise to the eukaryotic cell body and the proteobacteria went on to become the mitochondria [1, 2, 3]. Because of this, many physical processes that control eukaryotic cell division likely originate in archaea. In particular, ESCRT-III filaments, which drive cell division in a subgroup of archaea called TACK archaea, also catalyze the final step of cell division in many eukaryotes [4, 5].

Here we develop the first physical model to study the archaeal cell division by ESCRT-III filaments. In archaea, ESCRT-III proteins polymerise into at least two distinct filamentous rings which likely form a copolymer that is adsorbed on the cytoplasmic side of the cell membrane. The first filamenteous ring (called CdvB) serves as a template for the assembly of a contractile ring (made of CdvB1 and CdvB2 proteins), see Figure 1. Recently it has been shown that the contractile ring is only free to exert forces to reshape the membrane once the template CdvB ring has been removed [6]. As the membrane constricts, the contractile CdvB1/2 ESCRT-III filament disassembles.

**Figure 1.**
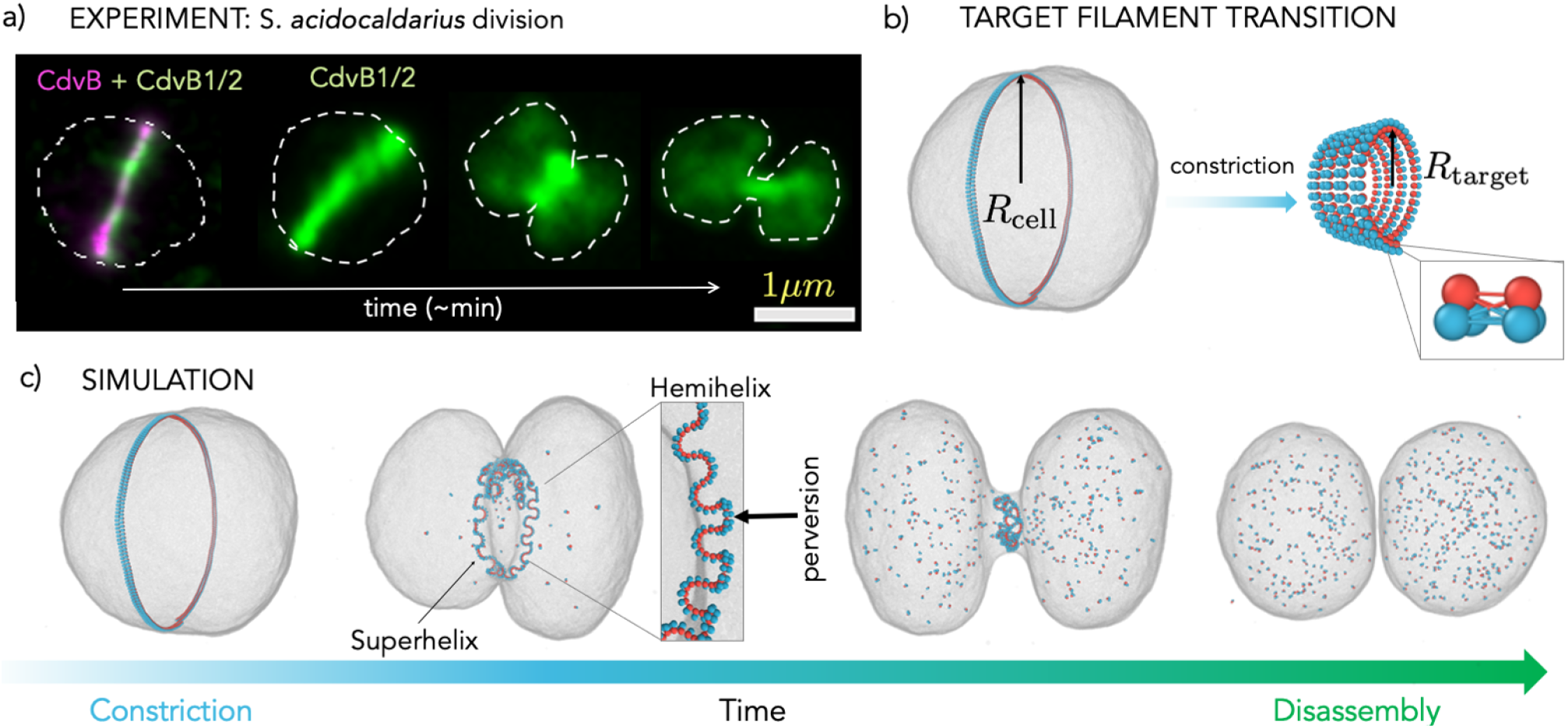
Computational model. a) Division of an archaeon *S. acidocaldarius*. The cell membrane is indicated with a dashed line, the template filament (CdvB) is fluorescently labelled in magenta, while the constricting ESCRT-III filament (CdvB1/2) is labeled in green. b) The initial ESCRT-III filament state (CdvB+CdvB1/2) is modelled as a single helical filament with a target radius equal to the cell radius *R*_cell_. The filament is attached to the inside of the fluid vesicle that represents the archaeal cell. To constrict, upon CdvB degradation, the ESCRT-III filament (CdvB1/2) reduces its target radius to *R*_target_, which results in a new target state of a tighter helix. The filament model itself consists of triplet subunits that are connected to each other via 9 bonds whose lengths control the filament curvature (see inset and SI). c) An example of a cell division simulation: the target radius of the filament is instantaneously decreased to 5% of the original cell radius. The filament is then disassembled from both ends with a rate 10^2^ × *v*_dis_ = 6.7*/τ* (*τ* is the MD unit of time). The filament first forms a superhelix which consists of multiple short helices of alternating chiralities (shown in the dashed box). As the superhelix contracts and disassembles, it pulls the membrane into a tight neck, which spontaneously breaks (Movie S1).

In our model we investigate different energy-driven protocols that can lead to the contraction and disassembly of an ESCRT-III ring in contact with a deformable cell. We quantify the rates, reliability, and symmetry of the resulting cell division processes. We then quantitatively compare the dynamics of cell division collected in simulations with that observed via live imaging of the archeon *Sulfolobus acidocaldarius*, the closest archaeal relative to eukaryotic cells that can be easily cultured in a laboratory. This comparison identifies a physical mechanism that matches all the available experimental data remarkably well.

Our study advances the understanding of the physical mechanisms governing cell division across evolution. In a broader sense, our simulations show how the different protocols that underpin the behaviour of a simple non-equilibrium nanomachine contribute to its function. Our results can hence be of help in designing protocols for synthetic nanomachinery, in particular in the context of building synthetic cells [7, 8, 9].

## Results

### A minimal model of cell division

In our model, the ESCRT-III filament (CdvB1/2) bound to the template (CdvB) is described as an elastic helical polymer made of two full turns whose radius of curvature matches the radius of the cell, *R*_cell_ (Figure 1b). The polymer is modelled via three beaded monomers that are connected by harmonic springs to each other (inset in Figure 1b). The equilibrium lengths of the springs determine the curvature of the polymer [10]. The archaeal cell is described as a vesicle made of a coarse-grained, one-particle-thick membrane (developed by Yuan et al. [11]), in which a single particle corresponds to a cluster of ~ 10 nm wide lipid molecules. These particles interact via an anisotropic pair potential that drives the self-assembly of fluid membranes with expected physiological bending rigidity. The outer side of the filament (coloured in blue beads in Figure 1b is attracted to the membrane via a generic Lennard-Jones attractive potential, while the inner side of the filament (red beads) only interacts with the membrane via volume exclusion. The system is evolved using molecular dynamics simulation with a Langevin thermostat (see Methods and Supplementary Information Section 1 for more details).

The removal of the template CdvB filament and subsequential constriction of the ESCRT-III filament (CdvB1/2) are modelled by shortening the equilibrium lengths of the bonds between the filament’s subunits, which increases the filament curvature, from a large radius of curvature *R*_cell_ to a small radius *R*_target_. Figure 1b shows the filament in its initial and target geometrical states. This change in target filament radius is expected to transform the filament from a wide ring into a tight helix and would drag the membrane with it. To test this idea, we instantaneously changed the target radius of the filament to 5% of the cell radius and let the system evolve. Since filament disassembly is needed for scission [6], as soon as the new equilibrium bond lengths have been imposed, we initiated the disassembly process by severing bonds between filament monomers sequentially from both ends of the helix at a specified rate *v*_dis_.

Interestingly, rather than adapting the target shape of a single tight helix, we find that the filament upon constriction instantly transforms into a collection of short tight helices of alternating chiralities separated by kinks called perversions (Figure 1c). Such a structure is called a hemihelix and it presents a local energy minimum in which the system gets trapped [12, 13, 14] (see SI Section 2). This shape arises due to non-homogeneous stresses in the filament, where the outer part of the filament is strained to a different extent relative to the inner part as the filament is changing its curvature. Hemihelical shapes, first documented by Darwin in plant tendrils [15], are ubiquitous in nature and have been reported in the development of the gut tube [16], in synthetic elastomers [17, 18], as well as in an everyday example of wrapping ribbon coiling by scissors [19]. Recently, this shape has also been imaged in nanoscale biopolymers such as mitotic chromosomes *in vivo* [20] and, importantly, for ESCRT-III and FtsZ filaments *in vitro* (see examples in Fig. S7, or e.g. Fig. 1B in Ref. [21], Fig. 5G-I in Ref. [22], or Fig. 6 in Ref. [23]).

When attached to the cell, the hemihelix itself acquires a superhelical shape to follow the cell curvature (Figure 1c). This effectively shortens the filament length and thereby pulls the membrane with it. Over time, the filament further shortens by disassembling, until it reaches the geometry of a single ring of a curvature 1*/R*_target_ enveloped by a tight membrane neck. As the helix disassembles, the membrane neck breaks, giving rise to two separate cells (Movie S1). To the best of our knowledge, this mechanism of constriction by effectively coiling a coil has not been considered before in cellular systems.

### Characterising the reliability and symmetry of cell division

We next investigate the functionality of such a division mechanism: how reliable it is and how good is it at partitioning protein content between the two daughters. Figure 2 shows the dependence of the probability of cell division on the amount of the filament curvature change, *R*_target_*/R*_cell_, and the rate of the filament disassembly, *v*_dis_. We see that division occurs only if both the filament curvature change and its disassembly rate are in the right regime. When the change in the filament curvature is not sufficient, the resulting large hemihelical loops barely shorten the filament and the filament does not have enough tension to constrict the membrane to a width sufficiently narrow to induce fission. Therefore, as the filament disassembles, the cell inflates back to its original size (square label in Figure 2). In the opposite regime, if the reduction in the filament curvature is too large, the large tension stored in the filament causes filament detachment from the membrane, preventing scission (circle in Figure 2).

**Figure 2.**
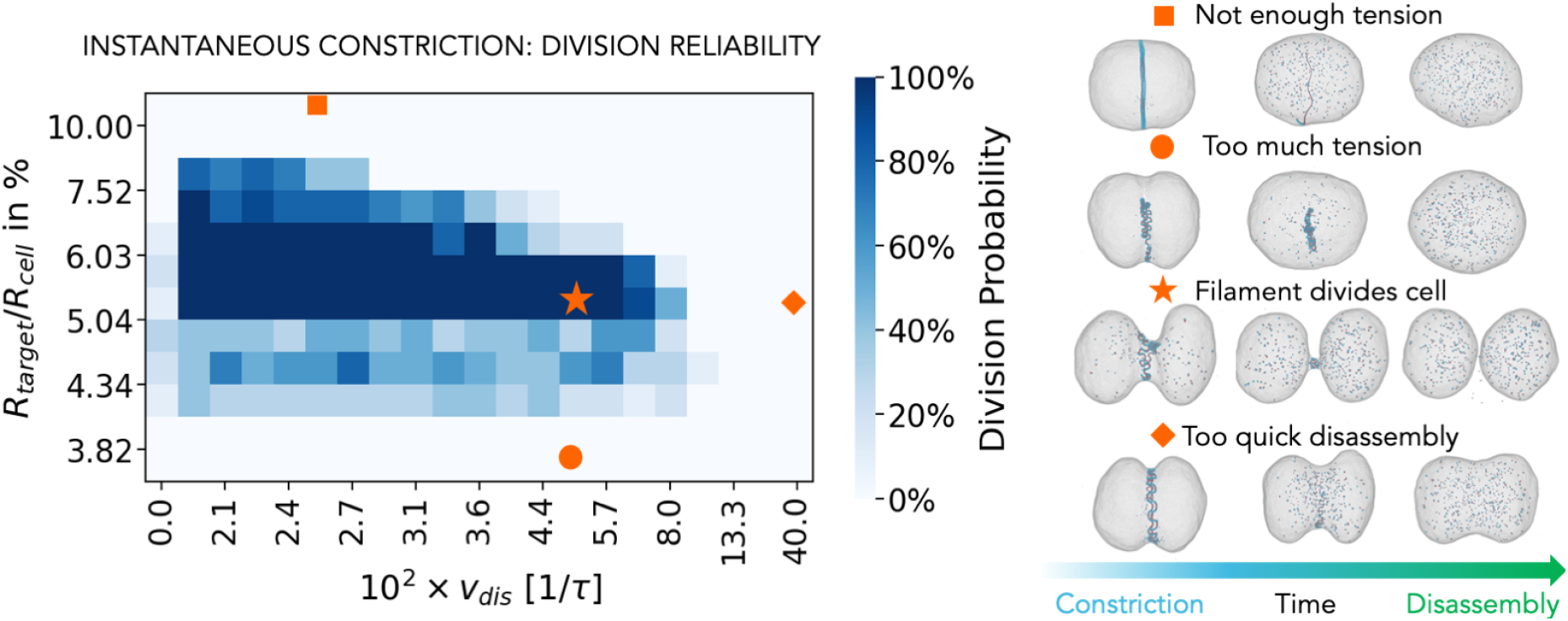
Reliability of the division. Left panel: The influence of the filament radius reduction, *R*_target_*/R*_cell_, and the rate of the filament disassembly, *v*_dis_, on the probability of cell division. The colour of each square represents the percentage of successful divisions out of 10 simulations performed with different random seeds. Right panel: The snapshots of the representative simulations.

In the intermediate regime, the filament has enough tension to form a sufficiently tight neck and is disassembled at a rate that then allows cell division (star in Figure 2). However, if the rate of disassembly is too high, the filament disassembles before the sufficiently small membrane bottleneck is reached, and division fails again (diamond in Figure 2). On the other hand, without disassembly (*v*_dis_ = 0), the filament generally blocks the membrane neck, preventing efficient cell division. Occasionally, the membrane neck can still break even in this case, leaving the daughter cells connected by the filament.

We also wanted to determine how the mechanism of division influences division symmetry. We therefore test how evenly the disassembled filament subunits are partitioned between the daughter cells for different curvature changes and disassembly rates, in the region of phase space in which division occurs. This evenness is quantified via parameter *E* = 2*N*_small_*/N*_total_, where *N*_small_ is the number of ESCRT-III subunits in the less filled daughter cell and *N*_total_ is the total number of ESCRT-III subunits.

As shown in Figure 3, for small filament curvature reductions, which is at the very boundary of the phase space in which division occurs (down-pointing triangle label), the filament does not constrict much and the division occurs because the membrane wraps over the exposed membrane attraction sites generated via local perversions in the helices formed. As a result, following division, one cell contains substantially more of the filament content than the other. This mechanism has a probability of yielding a successful division in only ~ 70% of cases (Figure 2). A similar effect occurs for strong radius reduction (up-pointing triangle label in Figure 3). Here the filament supercoils very quickly and the membrane wraps over one side of the coiled filament, leaving the filament in just one of the two cells. The result is again a less reliable and more uneven division. In the middle of the dividing region (star label in Figure 2 and diamond label in Figure 3), the protein partitions evenly between the daughter cells because the filament maintains a super-helical shape that is equally spread between the two cells. In this region the divison also occurs with ~ 100% probability.

**Figure 3.**
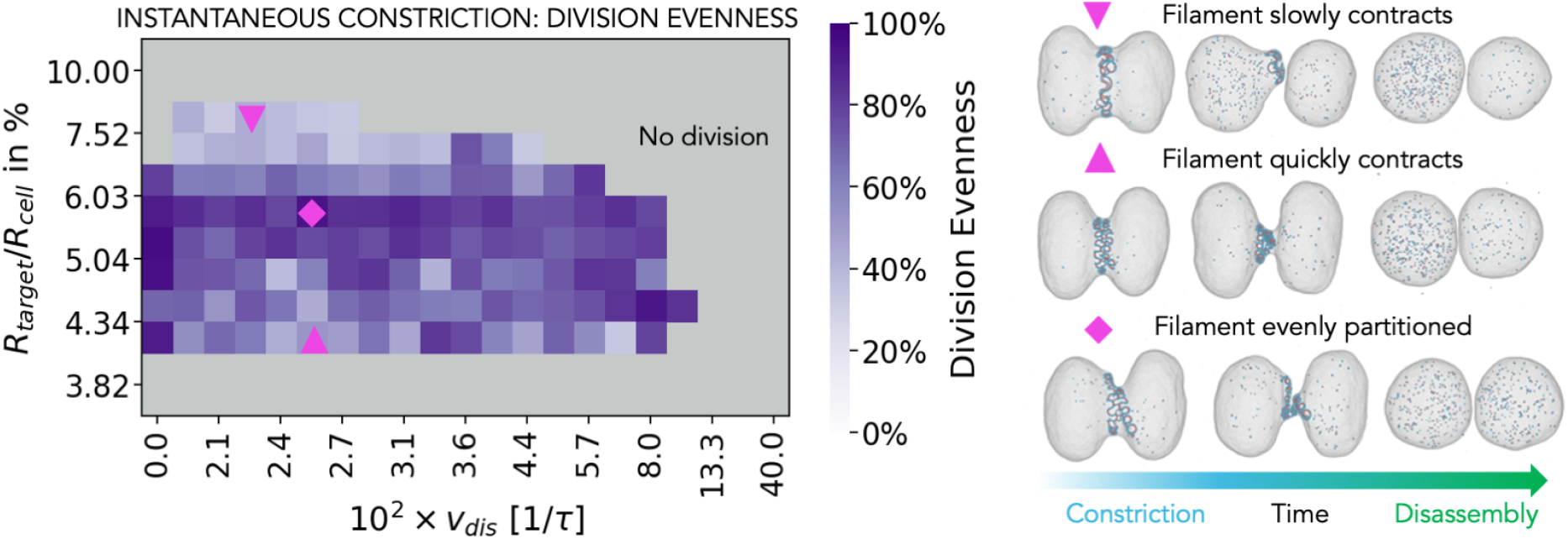
Symmetry of the division. Left panel: the influence of the amount of the instantaneous target radius reduction and the rate of the filament disassembly on the partitioning of ESCRT-III proteins between the daughter cells. The colour of each square represents average value of 10 simulations performed with different seeds. Right panel: Representative simulation snapshots.

### Dynamical curvature change protocols

The choice to induce instantaneous changes in filament curvature (Figure 4a) was inspired by experimental observations in *S. acidocaldarius*, where the template CdvB ring was found to be rapidly degraded, which was then assumed to trigger rapid curvature change of the constricting ring, akin to releasing of a loaded spring [6]. However, the local constriction rate is non-zero and is likely to be set by the local presence of an enzyme Vps4 ATPase, which is needed for division in *Sulfolobus*, as well as for force production of ESCRT-III filaments in eukaryotes [24, 25]. In fact, Pfitzner et al. [25] recently found that Vps4 modifies ESCRT-III filament composition in a step-wise manner, thereby changing the overall filament geometry, which in turn leads to membrane remodelling. This led us to investigate how different protocols of non-instantaneous dynamical filament constriction influence the outcome of the division process.

**Figure 4.**
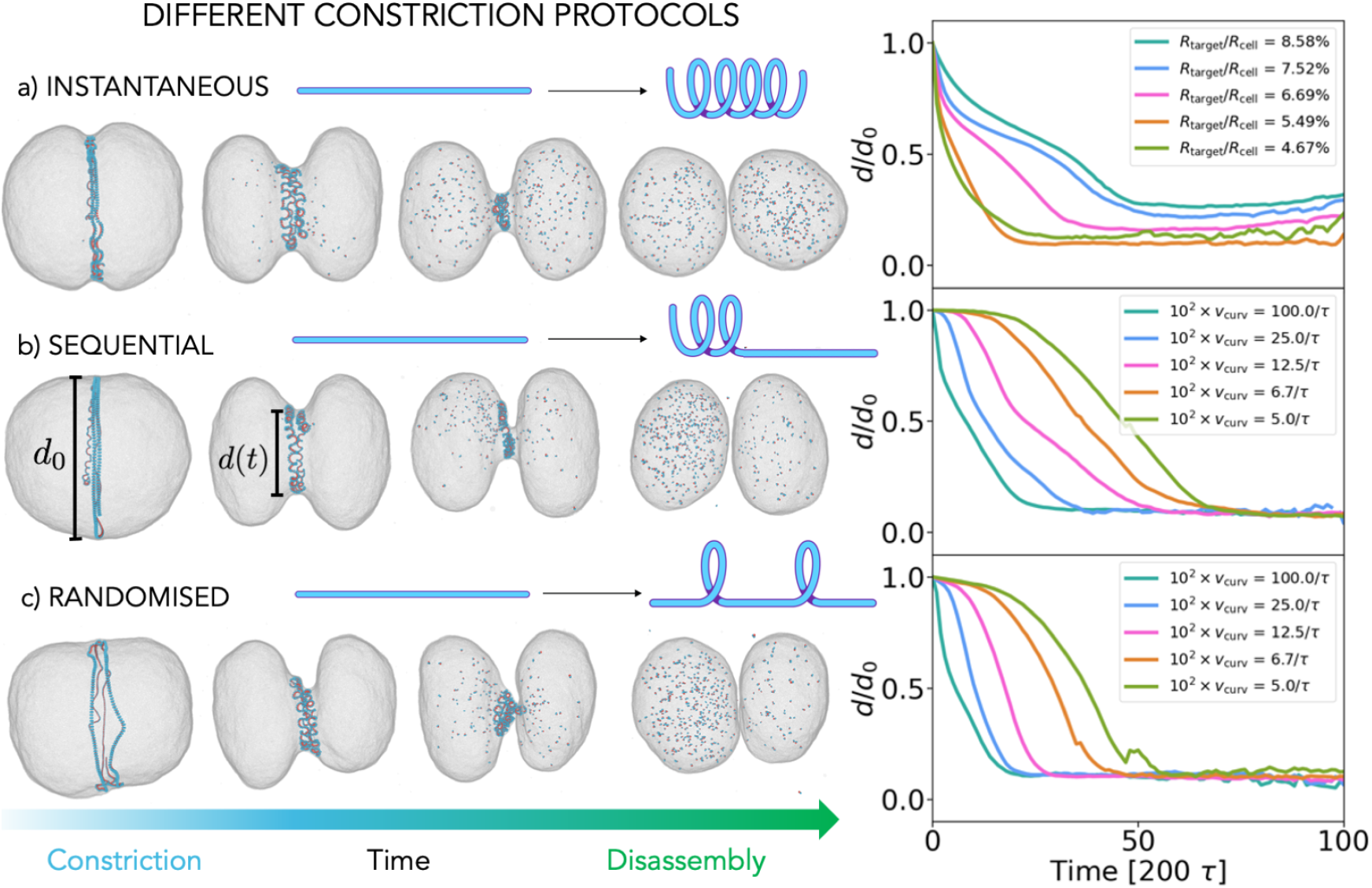
Different protocols for the curvature change. a) Instantaneous: The target filament radius changes globally throughout the filament to *R*_target_*/R*_cell_, as indicated in the legend, within a single timestep. b) Sequential: The filament curvature change starts at one end of the filament and propagates with a rate *v*_curv_ to the other end. Here shown for *R*_target_*/R*_cell_ = 5.5% c) Randomised: Random bonds along the filament constrict with a rate *v*_curv_. Here shown for *R*_target_*/R*_cell_ = 5.5%. In all the protocols: once the entire filament is transitioned, it is disassembled from both ends with a rate *v*_dis_. The value of *v*_dis_ does not influence the division curve (Figure S2). The panels on the right compare the normalised diameter of the cell at the midzone as a function of time. We only selected simulations that led to division and averaged over simulation seeds and different disassembly rates.

We test two additional protocols: i) filament curvature is changed in a sequential manner starting from one end of the filament in a process that spreads with a rate *v*_curv_ (Figure 4b) and ii) filament curvature is changed at random points along the filament with a rate *v*_curv_ (Figure 4c). In all the cases the filament disassembly starts after the new target curvature has been imposed throughout the whole filament. To compare these different mechanisms we track the cell diameter evolution in time for different protocols.

As can be seen in Figure 4a (right panel), following an instantaneous curvature change the cell diameter sharply decreases and then slowly plateaus before the division occurs. The smaller the target radius of the filament, the more energy is stored in the filament, and the faster the cell diameter reaches its equilibrium value. This behaviour is in a remarkably good agreement with the findings for the case of furrow behaviour in animal cell division by Turlier et al. [26].

The sequential and randomised protocols (Figure 4b and c) only follow the same qualitative curve if the curvature change is very fast, approaching the instantaneous limit; however if the curvature change is slower, the time evolution of the cell diameter follows a concave curve, where the furrow formation is significantly delayed, after which it follows an almost linear decrease before division occurs. The slower the constriction rate, the longer the delay in the furrow formation, which indicates that a certain critical portion of the filament has to be constricted in order to significantly deform the membrane. This delayed onset is less pronounced for the randomised protocol, since the filament is transformed at random points, the tension is distributed evenly across the midcell diameter ring. For the same reason, in the case of randomised constriction the diameter decreases faster than in the sequential protocol. Taken together, it is clear that the rate at which the filament geometry is changed greatly influences the relation between the diameter and time in both protocols: the curvature of the midcell diameter evolution changes from convex to concave as the rate of the curvature change decreases. Furthermore, we observe that the filament geometry itself depends on the rate of the curvature change: the slower the curvature change the less perversions the hemihelical filament contains (SI Section 2 and Figure S1), in agreement with previous studies of hemihelix formation in elastic materials [19].

Looking deeper into the difference between the sequential and random constriction protocols, we characterise the extent to which a robust cell division depends on the rate of the curvature change and the rate of the filament disassembly, fixing the amount of the curvature change to *R*_target_*/R*_cell_ = 5.5% (Figure 5a and b). In the case of the sequential curvature change, the cell only divides reliably (Figure 5a) and evenly (Figure S3a) for very fast curvature changes, which approach instantaneous constriction. In that regime the new curvature and associated tension spread quickly across the filament length, pulling the membrane with it (Movie S2). However, if the rate of the curvature change is decreased, the system enters a regime in which almost none of the cells divide. In this regime the tension is no longer spread evenly along the filament length and local filament loops have time to detach and equilibrate to their preferred curvature while the rest of the filament is still transitioning (Figure 5c and Movie S3). If the rate of the curvature change is however further decreased, the division becomes more likely again, because the detached local loops have time to reattach and pull the membrane with them.

**Figure 5.**
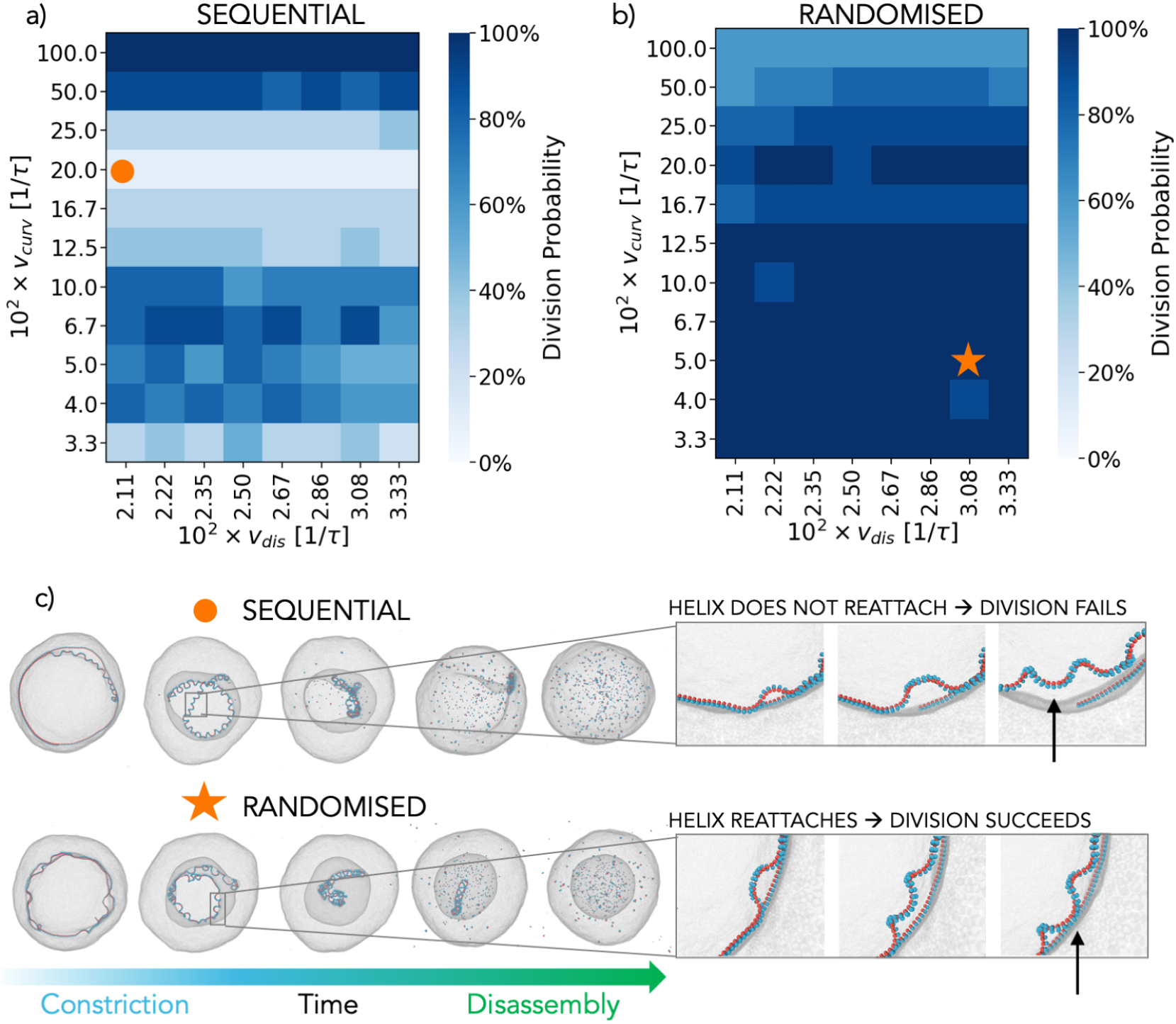
Reliability of division for non-instantaneous curvature change protocols. a) The influence of the filament constriction rate, *v*_curv_, and the filament disassembly rate, *v*_dis_, on the probability of cell division for the sequential (a) and randomised protocol (b). The colour of each square represents the amount of successful divisions out of 10 simulations performed with different seeds. c) The representative examples of a successful and unsuccessful division. If the local helices fail to reattach to the membrane when forming (indicated by an arrow in the upper panel), division fails. If the helices manage to reattach to the membrane (see arrow in the bottom panel), division suceeds. All simulations are performed using *R*_target_*/R*_cell_ = 5.5%, which divides cells with 100% reliability for the instantaneous protocol (Figure 2).

In the case of randomised curvature change protocol (Figure 5b, Movie S4), the curvature and the associated tension are distributed more evenly along the filament, no matter the rate, resulting in a more reliable and even (Figure S3b) division overall. Our model therefore predicts that cells divide more successfully and evenly if the constricting filament changes its curvature in a randomised fashion, rather than sequentially from one filament end.

### Quantitative comparison between simulation and experimental data

To identify which of the described constriction protocols best describes the ESCRT-III-driven archaeal cell division, we decided to compare the time evolution of the midcell diameter collected in simulations with similar measurements taken from live dividing *S. acidocaldarius* cells. To do so, we analysed cells imaged live using a membrane dye from Pulschen et al. [27], as shown in Figure 6a. This set-up allowed us to measure the intensity profile of the membrane along the division axis over time and to extract the value of the midcell (furrow) diameter evolution (Figure 6a, Figure S5-6). Since cells can have varying initial diameters and can take a varying time to divide, we rescale the midcell diameter with the initial cell diameter and the time axis with the total division time for each cell. We then align all the curves at the point of 50% of the cell diameter decrease. Satisfyingly, all the cells appear to follow the same general trajectory as they divide, which allows us to average the data into a single experimental curve (Figure 6b, see SI Section 6-7 for details). We transform our simulation measurements in the same way for a direct comparison with experimental data.

**Figure 6.**
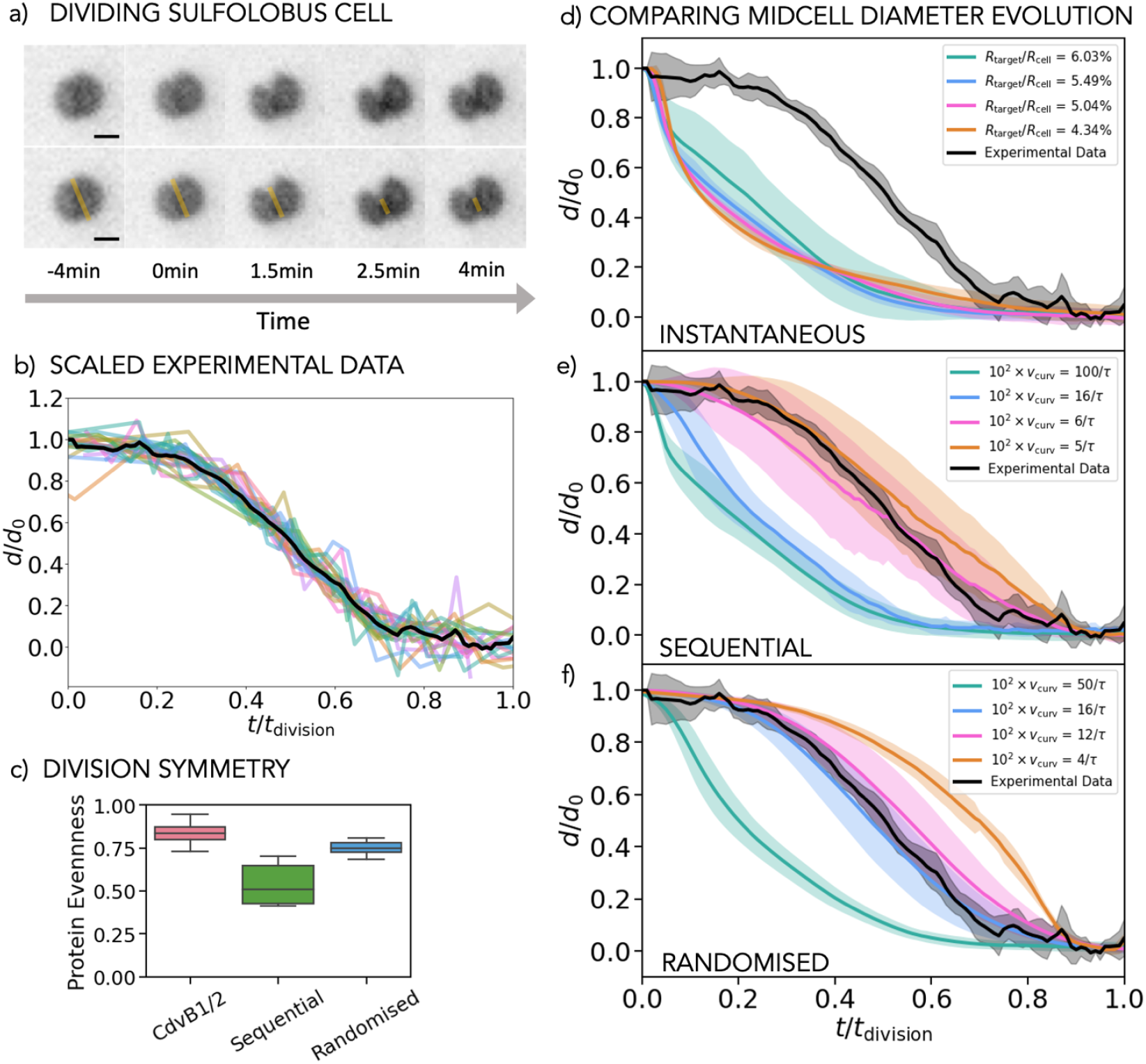
Comparison with live cell experiment. a) Time sequence of a dividing *S. acidocaldarius* cell. The bottom row shows the division axis along which we measure the membrane intensity profile to determine the time evolution of the midcell (furrow) diameter. Scale bar is 1*μm*. b) The evolution of the midcell diameter in time. The midcell diameter *d* is normalised by the initial cell diameter *d*_0_, while the time is normalised by the total division time *t*_*division*_. Coloured curves show the normalised measurements for individual cells (*N* = 23), and the black line shows the mean of all the experimentally measured curves. c) The average of the partitioning of the constricting filament proteins in the daughter cells in experiments (CdvB1/2) and in simulations for non-instantaneous curvature changes protocols that match the experimental curves for midcell evolution in time (10^2^*v*_curv_ = 5*/τ* for the sequential and 10^2^*v*_curv_ = 15*/τ* for the randomised protocol). d)-e) Show the normalised cell diameter evolution curves collected in simulations for different protocols (d: instantaneous; e: sequential; f: randomised) and compare it to the averaged experimental data (black curve). For the instantaneous protocol different amounts of filament constriction *R*_target_*/R*_cell_ are investigated. For the sequential and randomised protocols *R*_target_*/R*_cell_ = 5.5% and the constriction rate *v*_curv_ is varied, as shown. The disassembly rate does not influence the curves. All the simulation curves are averaged over at last 10 repetitions and the shading shows one standard deviation.

As can be seen in Figure 6d, for the instantaneous filament constriction, the behaviour of the furrow diameter in time has a very different qualitative shape than the experimental curve, irrespective of the amount of the curvature change in the filament. A possible reason for this discrepancy could be that our simulated cells are empty on the inside and do not resist the fast initial cell constriction when the target filament geometry is suddenly changed. To check the role of the cytoplasmic volume, we repeated the instantaneous constriction protocol measurements for cells that were filled with volume-excluded particles that mimic the cytoplasmic content. As Figure S4 shows, under these conditions the constriction curves do not change significantly; if division occurs the process is completely governed by the filament tension.

Using slower non-instantaneous filament curvature change protocols, the profiles develop initial and final plateaus and become more symmetric, resembling the experimental curve. Interestingly, the randomised protocol shows somewhat better agreement with the experimental curves, and for a range of simulation parameters, the normalised simulation and experimental curves match remarkably well (the blue and pink curves in Figure 6f). These best matching curves in the case of randomised curvature change also exhibit a reliable division (Figure 5b) and divide the protein evenly between the two daughter cells (Figure S3b).

For a further comparison between simulations and experiments, we also measured the evenness of filament protein (CdvB1/2) partitioning in daughter cells in experiments. Figure 6c shows the partitioning for CdvB1/2 proteins that we extracted from experiment, compared to the average evenness of partitioning in the case of the two best-matching curves in our simulated data (10^2^*v*_curv_ = 5*/τ* for the sequential and 10^2^*v*_curv_ = 16*/τ* for the randomised protocol). The experimental data shows that the filament proteins are divided fairly evenly between the daughter cells (Figure 6c), which agrees very well with the behaviour found in simulations for the randomised protocol. Taken together, our results indicate that a constriction protocol in which the filament curvature is non-instantaneously changed at random points throughout the filament length faithfully reproduces *Sulfolobus* cell division as measured using live cell imaging.

## Discussion

Here we develop a computational model for archaeal cell division via ESCRT-III filaments to study the physical mechanisms that underlie the division process. Our mesoscale model allows for a direct connection between the geometrical and mechanical properties of the filament and the resulting experimentally observable cellular behaviour. The model can hence be used as an *in silico* testing machine for division hypotheses that cannot be directly addressed in experiments.

One of the key questions about ESCRT-III’s ability to divide cells was how one system can constrict the membrane from a micrometre to just a few nanometres. Our model reveals the details of two important length-scales that are involved – the global curvature that arises due to the superhelical arrangement of a coiled filament and the local curvature of the individual coils. This supercoiling shortens the filament and guides the initial cell constriction, while the membrane wrapping around the individual tight coils creates a narrow neck that snaps after the filament has completely disassembled.

We find that the filament curvature changes that occur at random points along the filament provide a good model of the time evolution of the furrow diameter dividing *Sulfolobus* cells. Such a geometry change spreads the filament tension evenly along the filament, and hence yields the most reliable and symmetric division in the simulations. It can be imagined that it is also easiest for cells to implement this mechanism, as it does not rely on starting at a certain point within the filament.

Our model cannot determine which exact biochemical processes will create the randomised curvature change, but it is tempting to speculate that this is related to the ability of Vps4 ATPase enzyme to act in a nonprocessive way to extract the proteins of the CdvB template at random positions from the CdvB/B1/B2 copolymer. It should also be noted that, in our model the geometry changes and disassembly are implemented *ad hoc* and are uncoupled. It is, nevertheless, possible that the curvature changes and disassembly occur concomitantly, possibly through dependence of Vps4 activity on curvature or tension in the polymer. It is also possible that these events occur in a cooperative manner, giving rise to non-linear filament transformation. Since disassembly in our model does not influence the shape of the midcell diameter evolution curves, we cannot differentiate between more complex coupling scenarios of this type and, in the absence of further experimental data on the filament structure changes, we have opted for the simplest mechanism.

Our near-minimal model for cell division in this relatively simple archaeal cell system can also be of interest to those invested in generating synthetic cells, for which purposes the reconstitution of ESCRT-III filaments in vesicles is of substantial interest. Moreover, the analysis presented helps us to understand division of soft compartments by filaments in general, and forms the basis for studying more complex division mechanisms that evolved from this minimalistic archaeal division, such as severing of the midbody in the last step of eukaryotic division. In this light, under some conditions FtsZ has been seen to form similar hemihelices to those described here [23]. Finally, our model presents an excellent playground for testing specific non-equilibrium protocols behind bio-insipred nanomachines and their connection to the resulting nanomachine function.

## Acknowledgments

We acknowledge support from the Biotechnology and Biological Sciences Research Council (L.H.K.), EPSRC (A.E.H), UCL IPLS (T.Y and D. H.), Wellcome Trust (203276/Z/16/Z, A.P., S.C., R. H., B.B.), Volkswagen Foundation (Az 96727, A.P., B.B., A.Š.), MRC (MC CF1226, R.H., B.B., A.Š.), the ERC grant (”NEPA” 802960, A.Š.), the Royal Society (C.V.-H., A.Š.), the UK Materials and Molecular Modelling Hub for computational resources (EP/P020194/1).

## Author contributions

A.Š. and B.B. designed research. L.H.-K. and A.E.H. developed the computational model and performed simulations with help of C.V-H. and D.H.. T.Y., A.P., F.H and S.C. analysed experimental data. A.Š., B.B. and R.H. supervised research. All authors contributed to data analysis and paper writing.

## Methods

### Simulation

The simulation set-up – an elastic self-avoiding helical filament that is coupled to the inside of a spherical membrane, representing the archaeal cell – can be seen in Figure 1b. The filament helix has two full turns that consist of *N*_*total*_ = 480 subunits. The ratio between the helix width and diameter (6.5*σ/*105*σ ≈* 0.06) corresponds roughly to the experimentally measured ratio (0.1 *μ*m/1.25 *μ*m=0.08 [6]).

We simulate the filament using the ESCRT-III model we developed in Harker-Kirschneck et al. [10]. It is built of three-beaded rigid subunits that are bonded to each other via nine strong springs with spring constant *K*=600 k_B_*T* (inset in Figure 1b), where k_B_ is Boltzmann’s constant and *T* is temperature. The equilibrium lengths of the individual springs determine the target filament geometry (SI Section 1). The cell membrane is simulated by a coarse-grained, one-particle-thick model (developed by Yuan et al. [11]), in which a single particle corresponds to a lipid patch of ~ 10nm. These particles interact via an anisotropic pair potential that drives the self-assembly of fluid membranes with bending rigidity of 15 k_B_*T*. The blue beads of the filament are attracted to the membrane via a Lennard-Jones potential, while the red beads only interacts with the membrane via volume exclusion. A more detailed description of the simulation setup and all the interactions can be found in SI Section 1.

To change the filament curvature, the rest length of the bonds between the subunits is adjusted to the target curvature. In the case of non-instantaneous curvature change, the rate of the curvature change is determined by the number of bonds that are shortened per simulation timestep (0.01 *τ*), i.e. if we constrict the bonds between two subunits every *n*-th timestep, the rate is *v*_curv_ = 1/(*n*(0.01*τ*)). To simulate filament disassembly, every *m*-th simulation time step one subunit is removed from each end of the filament, making the disassembly rate *v*_dis_ = 2/(*m*(0.01*τ*)).

To measure the diameter *d* of the membrane at the cell midzone as a function of time, we collect all membrane particles that are located in a cuboid which is fixed at the centre of the simulation box and has a width of 1*σ* (the size of one membrane bead). The collected coordinates are then projected onto a plane and the resulting data is fitted by a circle using the Taubin method [28].

### Experiment

*Sulfolobus* cells were grown in BNS media and imaged using Nile Red for membrane staining, as described by Pulschen et al. [27]. This staining allowed us to measure the intensity profile of the membrane along the division axis over time (Figures S5 and S6.) The full width at half maximum (FWHM) of this profile was used as a measure of the midcell diameter of the dividing cells, a parameter that we can compare well to our simulation results (for details see SI Section 6). SI Section 7 describes the rescaling procedure of the data. It is important to add that for the last part of the midcell evolution curve the experimental uncertainty in measurement becomes larger due to the resolution limit, hence in all our results we focus on and discuss only the first half of the division curves.

## Data availability

All the simulation and experimental data, as well as simulation and analysis codes, are freely available upon request.

## References

[1] David A Baum and Buzz Baum. An inside-out origin for the eukaryotic cell. BMC biology, 12(1):1–22, 2014.

[2] Anja Spang, Jimmy H Saw, Steffen L Jørgensen, Katarzyna Zaremba-Niedzwiedzka, Joran Martijn, Anders E Lind, Roel Van Eijk, Christa Schleper, Lionel Guy, and Thijs JG Ettema. Complex archaea that bridge the gap between prokaryotes and eukaryotes. Nature, 521 (7551):173–179, 2015.

[3] Buzz Baum and David A Baum. The merger that made us. BMC biology, 18(1):1–4, 2020.

[4] Ann-Christin Lindås, Erik A Karlsson, Maria T Lindgren, Thijs JG Ettema, and Rolf Bernander. A unique cell division machinery in the archaea. Proceedings of the National Academy of Sciences, 105(48):18942–18946, 2008.

[5] Megan J Dobro, Rachel Y Samson, Zhiheng Yu, John McCullough, H Jane Ding, Parkson Lee-Gau Chong, Stephen D Bell, and Grant J Jensen. Electron cryotomography of escrt assemblies and dividing sulfolobus cells suggests that spiraling filaments are involved in membrane scission. Molecular biology of the cell, 24 (15):2319–2327, 2013.

[6] Gabriel Tarrason Risa, Fredrik Hurtig, Sian Bray, Anne E. Hafner, Lena Harker-Kirschneck, Peter Faull, Colin Davis, Dimitra Papatziamou, Delyan R. Mutavchiev, Catherine Fan, Leticia Meneguello, Andre Arashiro Pulschen, Gautam Dey, Siân Culley, Mairi Kilkenny, Diorge P. Souza, Luca Pellegrini, Robertus A. M. de Bruin, Ricardo Henriques, Ambrosius P. Snijders, AndelaŠ arić, Ann-Christin Lindås, Nicholas P. Robinson, and Buzz Baum. The proteasome controls escrt-iii–mediated cell division in an archaeon. Science, 369(6504), 2020. ISSN 0036-8075. doi: 10.1126/science.aaz2532. URL https://science.sciencemag.org/content/369/6504/eaaz2532.

[7] Andrew Booth, Christopher J Marklew, Barbara Ciani, and Paul A Beales. In vitro membrane remodeling by escrt is regulated by negative feedback from membrane tension. Iscience, 15:173–184, 2019.

[8] Yuval Elani, Tatiana Trantidou, Douglas Wylie, Linda Dekker, Karen Polizzi, Robert V Law, and Oscar Ces. Constructing vesicle-based artificial cells with embedded living cells as organelle-like modules. Scientific reports, 8(1):1–8, 2018.

[9] Siddharth Deshpande and Cees Dekker. On-chip microfluidic production of cell-sized liposomes. Nature protocols, 13(5):856, 2018.

[10] Lena Harker-Kirschneck, Buzz Baum, and Andela Šarić. Changes in escrt-iii filament geometry drive membrane remodelling and fission in silico. BMC Biology, 17(1):82, 2019.

[11] Hongyan Yuan, Changjin Huang, Ju Li, George Lykotrafitis, and Sulin Zhang. One-particle-thick, solvent-free, coarse-grained model for biological and biomimetic fluid membranes. Phys. Rev. E, 82: 011905, Jul 2010. doi: 10.1103/PhysRevE.82.011905. URL https://link.aps.org/doi/10.1103/PhysRevE.82.011905.

[12] Jiangshui Huang, Jia Liu, Benedikt Kroll, Katia Bertoldi, and David R. Clarke. Spontaneous and deterministic three-dimensional curling of pre-strained elastomeric bi-strips. Soft Matter, 8:6291–6300, 2012. doi: 10.1039/C2SM25278C.

[13] Jia Liu, Jiangshui Huang, Tianxiang Su, Katia Bertoldi, and David R Clarke. Structural transition from helices to hemihelices. PloS one, 9(4):e93183, 2014.

[14] Shuangping Liu, Zhenwei Yao, Kevin Chiou, Samuel I Stupp, and Monica Olvera De La Cruz. Emergent perversions in the buckling of heterogeneous elastic strips. Proceedings of the National Academy of Sciences, 113 (26):7100–7105, 2016.

[15] Piotr Pieranski, Justyna Baranska, and Arne Skjeltorp. Tendril perversion—a physical implication of the topological conservation law. European Journal of Physics, 25:613, 06 2004. doi: 10.1088/0143-0807/25/5/004.

[16] Thierry Savin, Natasza A. Kurpios, Amy E. Shyer, Patricia Florescu, Haiyi Liang, L. Mahadevan, and Clifford J. Tabin. On the growth and form of the gut. Nature, 476 (7358):57–62, 2011.

[17] Pedro ES Silva, Joao L Trigueiros, Ana C Trindade, Ricardo Simoes, Ricardo G Dias, Maria Helena Godinho, and Fernao Vistulo De Abreu. Perversions with a twist. Scientific reports, 6(1):1–8, 2016.

[18] PES Silva, F Vistulo De Abreu, and MH Godinho. Shaping helical electrospun filaments: a review. Soft Matter, 13(38):6678–6688, 2017.

[19] Chris Prior, Julien Moussou, Buddhapriya Chakrabarti, Oliver E Jensen, and Anne Juel. Ribbon curling via stress relaxation in thin polymer films. Proceedings of the National Academy of Sciences, 113(7):1719–1724, 2016.

[20] Lingluo Chu, Zhangyi Liang, Maria V Mukhina, Jay K Fisher, John W Hutchinson, and Nancy E Kleckner. One-dimensional spatial patterning along mitotic chromosomes: A mechanical basis for macroscopic morphogenesis. Proceedings of the National Academy of Sciences, 117(43):26749–26755, 2020.

[21] Joachim Moser Von Filseck, Luca Barberi, Nathaniel Talledge, Isabel E Johnson, Adam Frost, Martin Lenz, and Aurélien Roux. Anisotropic escrt-iii architecture governs helical membrane tube formation. Nature communications, 11(1):1–9, 2020.

[22] William Mike Henne, Nicholas J. Buchkovich, Yingying Zhao, and Scott D. Emr. The endosomal sorting complex escrt-ii mediates the assembly and architecture of escrt-iii helices. Cell, 151(2):356–371, 2018/11/28 2012.

[23] Erin D Goley, Natalie A Dye, John N Werner, Zemer Gitai, and Lucy Shapiro. Imaging-based identification of a critical regulator of ftsz protofilament curvature in caulobacter. Molecular cell, 39(6):975–987, 2010.

[24] Johannes Schöneberg, Mark Remec Pavlin, Shannon Yan, Maurizio Righini, Il-Hyung Lee, Lars-Anders Carlson, Amir Houshang Bahrami, Daniel H Goldman, Xuefeng Ren, Gerhard Hummer, et al. Atp-dependent force generation and membrane scission by escrt-iii and vps4. Science, 362(6421):1423–1428, 2018.

[25] Anna-Katharina Pfitzner, Vincent Mercier, Xiuyun Jiang, Joachim Moser von Filseck, Buzz Baum, Andela Šarić, and Aurélien Roux. An escrt-iii polymerization sequence drives membrane deformation and fission. Cell, 182(5): 1140–1155, 2020.

[26] Hervé Turlier, Basile Audoly, Jacques Prost, and Jean-François Joanny. Furrow constriction in animal cell cytokinesis. Biophysical journal, 106(1):114–123, 01 2014.

[27] Andre Arashiro Pulschen, Delyan R Mutavchiev, Siân Culley, Kim Nadine Sebastian, Jacques Roubinet, Marc Roubinet, Gabriel Tarrason Risa, Marleen van Wolferen, Chantal Roubinet, Uwe Schmidt, et al. Live imaging of a hyperthermophilic archaeon reveals distinct roles for two escrt-iii homologs in ensuring a robust and symmetric division. Current Biology, 30(14):2852–2859, 2020.

[28] N. Chernov. Circular and Linear Regression: Fitting Circles and Lines by Least Squares. Chapman & Hall/CRC Monographs on Statistics & Applied Probability. Taylor & Francis, 2010. ISBN 9781439835906. URL https://books.google.co.uk/books?id=GaKTQgAACAAJ.

