## Supplementary Information for "Physical mechanisms of ESCRT-III-driven cell division in archaea"

This section contains: (1) details on the simulation set-up, (2) additional simulations on the relationship between the rate of filament constriction and the filament geometry, (3) measurements on the dependence of the time evolution of midcell diameter on the rate of the filament disassembly for instantaneous constriction, (4) measurements on the division evenness for the sequential and randomised protocols, (5) additional simulations and measurements for the case when the cell is filled with cytoplasmic particles, (6) analysis of the cell volume and area changes during division, (7) experimental details on cell imaging, (8) details of the fitting of the time evolution of the midcell diameter data, and (9) information on Supporting Movies S1-4, (9) examples of hemihelices observed experimentally in yeast ESCRT-III filaments.

### 1. Simulation set-up

The cell membrane is modelled using the coarse-grained, solvent free, one-particle-thick membrane model by Yuan et al. [1]. Following the original paper, the membrane beads have a diameter $\sigma$, where $\sigma$ is also the MD unit of length, and the inter-bead interaction parameters are set to: $\varepsilon_{\mathrm{memb}}$=4.34 $\mathrm{k_B}T$, $\xi$=4, $\mu$=3, and $r_{\mathrm{cut}}$=1.12 $\sigma$. These parameters reproduce a fluid deformable membrane of a bending rigidity of 15 $\mathrm{k_B}T$, which is in the physiological regime. The membrane includes 48002 particles and forms a vesicle of radius $R_{\mathrm{cell}} = 52.464\ \sigma$.

The ESCRT-III filament in the model consists of 3 beaded subunits, each of diameter $\sigma$, which form rigid bodies (see inset in Figure 1b). The beads of neighbouring subunits are connected by nine harmonic bonds, whose spring constant is set to $600 k_B T$. The bond lengths are set to result in an intrinsic filament curvature and the filament curvature can be adjusted by varying the bond lengths. This curvature is the same throughout the filament, hence the relaxed state of the filament is a ring of a radius $R_{\mathrm{target}}$. Since the filament subunits cannot overlap with each other, if the filament consists of more than one turn, this resulting relaxed geometry will be a helix instead of a ring.

The blue beads in the filament interact with membrane beads that are at distance $r_{ij}$ via a cut-and-shifted Lennard-Jones potential: $E_{ij} = 4\varepsilon\left((\sigma/r_{ij})^{12} - (\sigma/r_{ij})^{6}\right)$, $r_{ij} < r_c$, with $\varepsilon$=4 $\mathrm{k_B}T$ and the cut-off-distance $r_c$=1.46 $\sigma$. In contrast, the red beads of the filament and the membrane particles, as well as the beads of the filament with each other, only interact via volume exclusion, implemented via Lennard-Jones potential of $\varepsilon$=2 $\mathrm{k_B}T$, cut and shifted at the minimum of the potential.

We perform molecular dynamics (MD) simulations using the molecular dynamics package LAMMPS [2] to integrate the equations of motion with periodic boundary conditions in an $N_{particles}V_{box}E_{system}$ ensemble, where $N_{particles}$ is the total number of particles in the box of a volume $V_{box}$ and $E_{system}$ is the total energy of the system. The simulation box is a cube with a fixed edge length of $L_{\mathrm{box}} = 200\,\sigma$. All particles in the system experience Langevin dynamics with the friction coefficient set to unity, $\gamma$=$m/\tau$, where $m$ is the particle mass (set to unity for all particles) and $\tau$ is the MD unit of time. The MD time-step was chosen to be $0.01\tau$. At the beginning of every simulation, we first equilibrate the vesicle on its own, and then the filament in contact with the vesicle, where the target radius of the filament equals that of the cell. At the end of this equilibration stage the cells look like in the first snapshots in Figure 1c. The simulation results are visualized using OVITO [3], an open-source analysis and visualization tool.

### 2. Constriction rate vs filament geometry

In order to study the effect of the rate of the curvature change on the filament geometry, we repeat the constriction simulation (as in Figure 1c) but without a membrane. Figure S1a shows snapshots from such a constriction to 5% of the original cell radius. As in the equivalent simulation that includes a membrane (compare Figure 1c), the filament constricts by forming helical-like structures that are frequently interrupted by changes of chirality, known as perversions [4].

We then studied the influence of the constriction rate $v_{\mathrm{curv}}$ on the geometry of the filament. Figure S1b shows the number of perversions that occurred in the equilibrated filament as a function of $v_{\mathrm{curv}}$, along with as the total energy of the filament. A faster constriction leads to the formation of more perversions, which in turn leaves the filament in a more frustrated state, indicated by the rise in energy of the filament [4].

### 3. Midcell diameter dependence on disassembly rate

Constricting the filament can drive the formation of small bottlenecks, however, disassembly is necessary to achieve

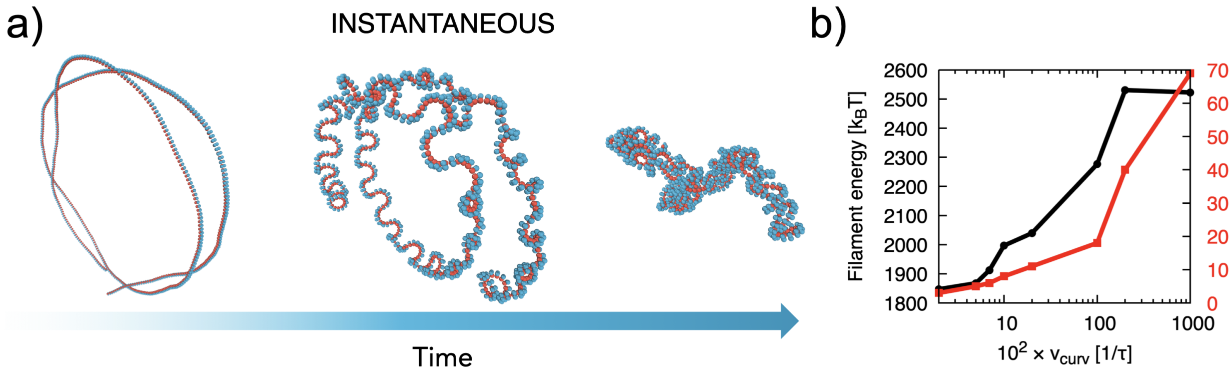


**Figure S1.** a) Reducing the target radius to 5% of the original cell radius for a filament that is not attached to a membrane using the instantaneous protocol. b) Final value of the filament energy (black line) and the corresponding number of helical perversions that developed (red line) as a function of the constriction rate $v_{curv}$.

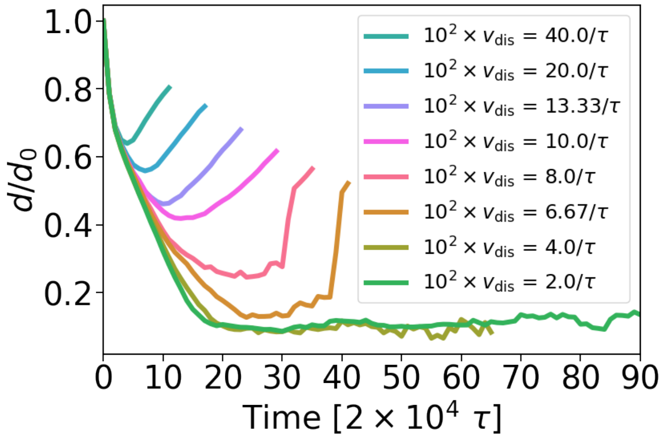


**Figure S2.** Midcell diameter evolution after constricting the filament instantaneously by $R_{target}/R_{cell} = 5.5\%$ and varying the disassembly rate $v_{dis}$. Each curve is the mean of 10 random seeds. Note that disassembly must be slow ($10^2 v_{dis} < 6.67\tau$) for divisions to complete.

robust cell division. The disassembly rate $v_{dis}$ has to be well adjusted to the time it takes for the bottleneck to establish. In Figure S2, we plot the cell diameter as a function of time for $R_{target}/R_{cell} = 5.5\%$ and various disassembly rate $v_{dis}$ following the instantaneous constriction protocol.

We can see that all lines initially follow the same general curve, but if the filament disassembles before it can form a sufficiently small bottleneck, the cell membrane recovers its initial spherical shape and the diameter increases again. For disassembly rates slower than $10^2 v_{dis} = 6.67\tau$, the bottleneck can be maintained long enough for the cells to divide. If we only look at cells that divide, the disassembly rate has no influence on the shape of the midcell evolution curve, hence we can use these simulations for additional statistics.

### 4. The division evenness for the sequential and randomised protocol

Figure S3 shows how evenly the cells divided on average (using 10 different seeds per square), depending on the disassembly rate $v_{dis}$ and the rate at which the filament contracts $v_{curv}$ for non-instantaneous constriction protocols. Division symmetry is measured by assessing how much of the ESCRT-III polymer ends up in the two daughter cells. For the sequential protocol the division is generally not very even, with the even division occurring only for the fast constriction rates (Figure S3a). The randomised regime division yields very even division, regardless of the constriction rate or the disassembly rate (Figure S3b).

### 5. The effect of cytoplasmic volume

The simulation set-up in the main paper does not account for fact that the cell is filled with cytoplasmic content, where 20-30% of its volume is occupied by proteins [5]. In order to investigate the presence of non-compressible cytoplasmic content on the dynamics of cell division, we place volume-excluded particles inside the simulated cell. The volume exclusion ins implemented via Lennard-Jones potential of $\varepsilon$=2 $k_B T$, cut and shifted at the potential minimum.

The hexagonal close packing arrangement was used to place the cytoplasmic particles inside the cells at high packing

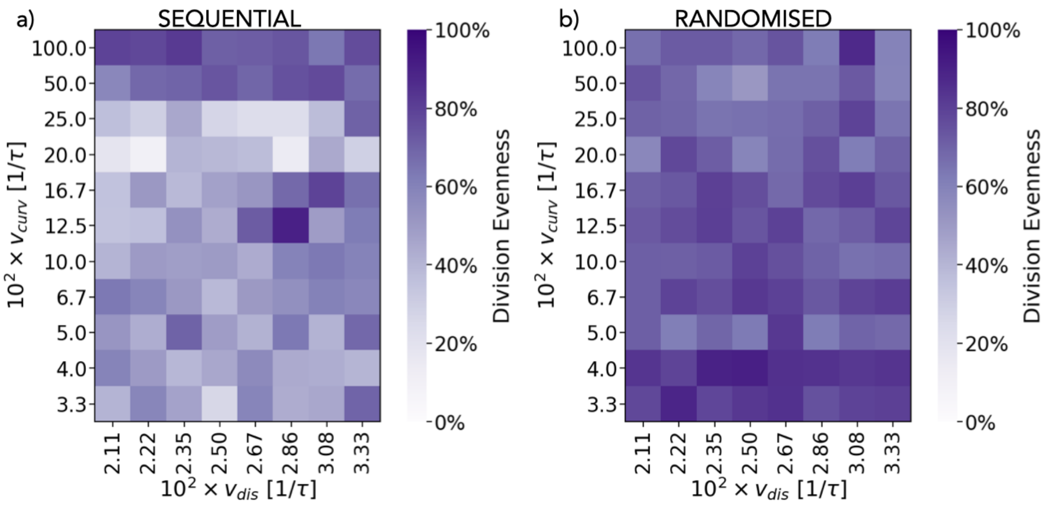


**Figure S3.** a) The influence of the constriction rate $v_{\mathrm{curv}}$ and the rate at which the filament is disassembled $v_{\mathrm{dis}}$ on how evenly the cells divide following the sequential protocol. The colour represents the division evenness $E$, as defined in the main text. If $E = 100\%$ the daughter cells contain the exact same amount of filament subunits. Each square shows the average Evenness $E$ out of 10 simulations performed with different seeds. b) The same as in a) but following the randomised regime. Here the cells all divide very evenly regardless of the parameters we chose.

fraction, as shown in Fig. S4a. We then explored the influence of the packing fraction (the percentage of volume inside the cell taken up with cytoplasmic particles), as well as the cytoplasmic particle diameter $\sigma_{\mathrm{cyto}}$, on cell division. Simulations were run for ten different seeds at packing fractions 10-40%, $\sigma_{cyto} = 5,6,7,8\sigma$ and for the three different curvature change protocols. We fix the amount by which we reduce the target radius of the filament to $R_{\mathrm{target}}/R_{\mathrm{cell}} = 6\%$ and the rate at which the filament disassembles to $10^2 v_{\mathrm{dis}} = 3.3/\tau$. The division probability depending on the packing fraction and the cytoplasmic particle diameter is represented via the colour of each square in Figure S4c. The number of cytoplasmic particles in each simulation is also included in the diagram.

Fig. S4c shows that the greater the number of cytoplasmic particles, the less likely that cell division is successful. This result is to be expected as the presence of particles within the simulated cell causes internal pressure that acts against the constricting force of the filament protein that drives cell division. Figure S4d shows the time evolution of the average midcell diameter (over 10 seeds) for varying packing fractions using the randomised protocol. For all packing fractions, the filament was constricted to 6% of its original cell radius and the cytoplasmic particle radius was fixed at $\sigma_{cyto} = 8$, because it is the configuration with the greatest amount of variation in cell division success.

The greater the packing of the cell, the slower the rate of the constriction. At very high packing fraction the division fails. Importantly, if the division succeeds (for the curves $0-25\%$ packing), the midcell evolution curve always follows the same general shape. Hence, the difference between the experimental and simulation curve in Figure 6d likely cannot be explained by the lack of the resistance of the cytoplasmic volume in the initial model.

### 6. Measurements of the midcell diameter

The microscopy field of view covered an area of $\sim 150\mu m \times 150\mu m$, displaying multiple cells at once. We used a machine learning algorithm to detect any dividing cells and cropped them from the master image stack (using ImageJ) to a more appropriate scale as can be seen in Figure S5.

The resolution of movies of dividing *S. acidocaldarius* were limited by the small size of the cells and the resolution of the microscope. The Full-Width Half-Maximum (FWHM) of the point-spread function of the microscope for this experiment was estimated to 632 nm. It is therefore not straightforward to determine the edges of the cell. The FWHM of an intensity profile can be used to describe the measurement of the width of an imaged object when the edges of the image are not sharp [6]. The intensity profile of the midcell diameter can be extracted using ImageJ's line tool as shown in Figure S7a.

We then fitted the intensity profiles via a Gaussian, and calculated FWHM of the fit can be calculated via

$$FWHM = 2\sqrt{2ln2}\sigma_G \approx 2.355\sigma_G. \quad \text{(S1)}$$

where $\sigma_G$ is the standard deviation of the fitted Gaussian.

Some images of the cells have two intensity peaks corresponding to two Gaussians. The two Gaussians are present in the early stages of cell division, as shown in the left panel of Figure S7b. This is due to the overlapping membranes of the daughter cells that create two localised areas of increased intensity. As the cell divides, the two areas move closer together until the two intensity peaks form one. Hence, initial intensity measurements are plotted to fit two Gaussian curves, and the later measurements are plotted to fit one Gaussian.

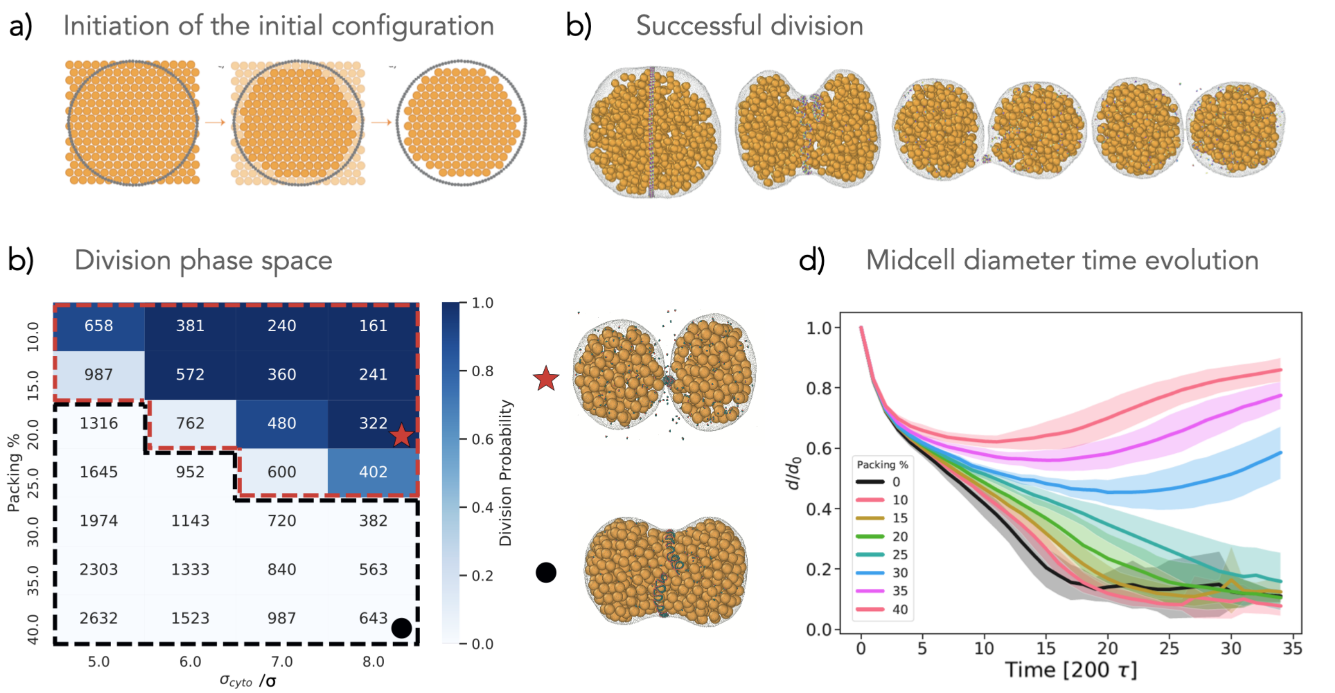


**Figure S4.** Investigating the role of the cytoplasmic volume on the division. a) The protocol of how cytoplasmic particles are initialised in the cell model. b) A representative example of a successful division. Here the diameter of the cytoplasmic particle is $\sigma_{cyto} = 8\sigma$ and packing fraction is 20%. c) The probability of successful cell division depending on the packing fraction and the diameter of the cytoplasm particles, $\sigma_{cyto}$. The number inside each cell represents the number of cytoplasm particles in the cell. b) Midcell diameter against time dependent on the packing of cytoplasmic particles, where $\sigma_{cyto} = 8\sigma$.

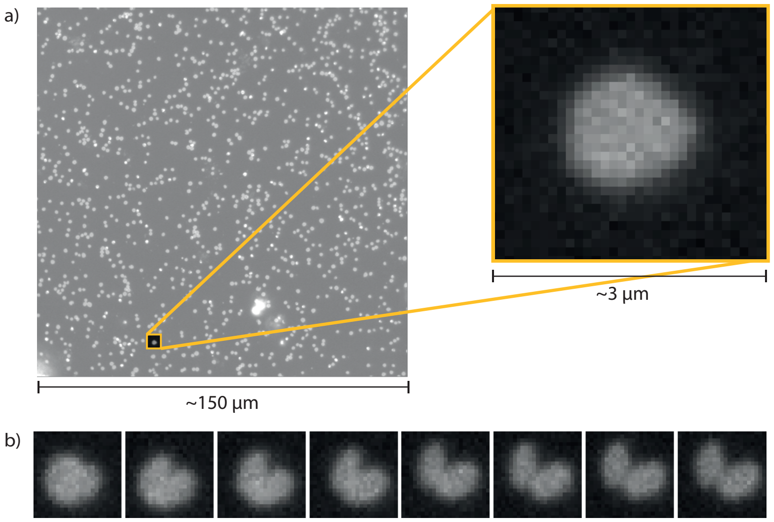


**Figure S5.** a) A dividing cell cropped out of a time-lapse film of MW001 *S. acidocaldarius* cells using ImageJ software. b) A montage of images showing how the cropped cell divides over time, with one image taken every minute.

If two peaks were present, two Gaussians were fitted to the profile instead of one. These fits were background-subtracted such that their offsets were zero (Figure S7b, middle panel), and then combined to create a unified curve (Figure S7b, right panel). This process was repeated for each frame of the data where the profile was poorly represented by a single Gaussian. The combined double-Gaussian fit does not have a single well-defined $\sigma_G$. In this case, the FWHM was measured as the width of the combined fit profile corresponding to x-values at half the intensity of the higher-amplitude peak.

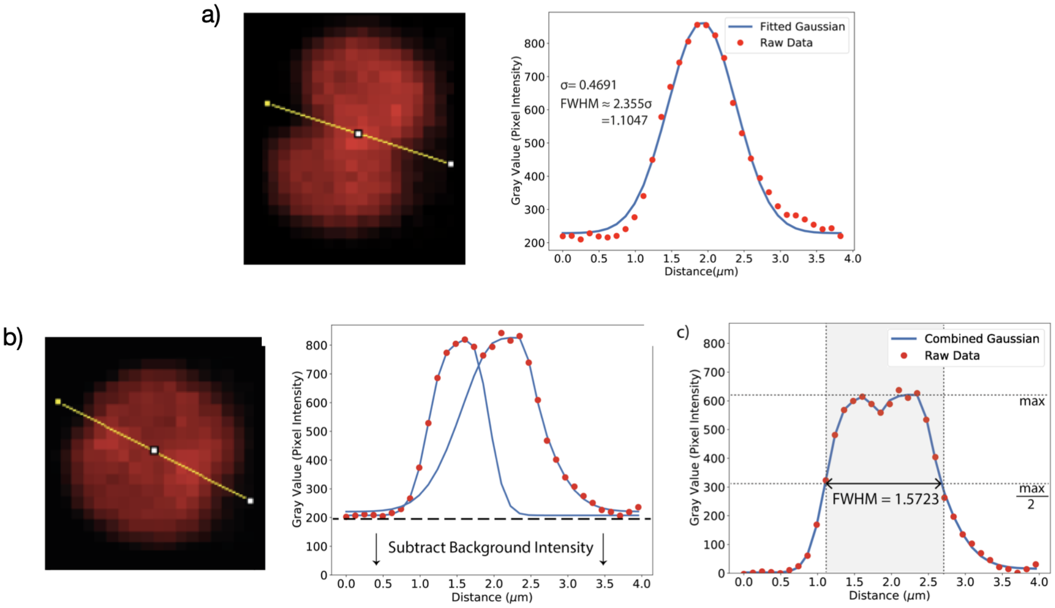


**Figure S6.** a) Left panel: The intensity over the midcell axis of a cell is measured using the line tool in ImageJ. Right panel: The intensity is plotted against distance and fitted with a Gaussian curve. The FWHM of the Gaussian is calculated by multiplying the standard deviation of the curve by $2\sqrt{2ln2}$. b) In the early stages of division, cells appear to have two intensity peaks on the midaxis of the cell due to the overlapping of daughter cells. Left panel: The intensity over the midcell axis of a cell is measured using the line tool in ImageJ. Middle panel: The intensity is plotted against distance and fitted with two Gaussian curves, then zeroed by subtracting the background intensity. Right panel: The two Gaussians are then joined, and the FWHM is calculated from the combined Gaussian.

### 7. Midcell diameter evolution rescaling

The shape of the curves describing the midcell diameter of the dividing cells over time resemble a sigmoidal function, but can be quite asymmetric if the filament curvature changes quickly. We hence fitted them using this generalised logistic function that allows the curves to be both asymmetric and symmetric:

$$Y(t) = A + \frac{K - A}{(1 + e^{-B(t-M)})^{1/v}} \tag{S2}$$

with $A$ being the lower and $K$ the upper asymptote. $B$ is the growth rate and $v$ determines how asymmetric the curve gets. $M$ can be considered the starting time at which the curve turns: $Y(t = M) = A + \frac{K - A}{2^{1/v}}$. To improve the fit quality, we chose $v = 1$ for the experimental fits, as they are very symmetric. For the simulated curves, we chose the upper asymptote $K = 1$, since we know they all start at the same original diameter.

We then fit the diameter over time (for all random seeds) using this function and extract the fitting parameters to scale the curve in the x and y direction. Scaling in the y-direction is simple, as we only need to subtract the minimum asymtopte value from the data points and then divide them by the difference between the upper and lower asymptote: $\text{Diameter}_{\text{scaled}} = (\text{Diameter} - A)/(K - A)$. Now the y-axis is given in % of original diameter, rather than the diameter in $\mu m$. Next we scale the x-axis from time in min to % of completed division. To do this we measure the time points at which the fits reach 99% ($T_{\text{start}}$) and 1% ($T_{\text{end}}$) of their maximal values respectively. We then subtract the starting time from all x-values and divide by the different between the starting end ending time, which is the amount of time the cell took to divide: $\text{Time}_{\text{scaled}} = (\text{Time} - T_{\text{start}})/(T_{\text{start}} - T_{\text{end}})$. Once scaled, we interpolate the midcell diameter evolution curves for all simulation seeds and calculate their mean and standard deviation, which is then displayed in Figure 6 d)-f).

### 8. Supporting Movies

**Movie S1**: Example of a successful cell division simulation following the instantaneous constriction protocol. Here $R_{\text{target}}/R_{\text{cell}} = 5\%$, $10^2 \times v_{\text{dis}} = 6.7/\tau$.
**Movie S2**: Example of a successful cell division simulation following the sequential constriction protocol for a fast curvature change rate, $10^2 \times v_{\text{curv}} = 100.00/\tau$. $R_{\text{target}}/R_{\text{cell}} = 5\%$ and $10^2 \times v_{\text{dis}} = 2.11/\tau$.
**Movie S3**: Example of a failed cell division simulation following the sequential constriction protocol performed at a slow rate, $10^2 \times v_{\text{curv}} = 20.00/\tau$. $R_{\text{target}}/R_{\text{cell}} = 5\%$ and $10^2 \times v_{\text{dis}} = 2.11/\tau$.
**Movie S4**: Example of a successful cell division following the randomised constriction protocol at a moderate rate, $10^2 \times v_{\text{curv}} = 5.00/\tau$. $R_{\text{target}}/R_{\text{cell}} = 5\%$ , $10^2 \times v_{\text{dis}} = 3.08/\tau$.

### 9. Hemihelices in ESCRT-III filaments

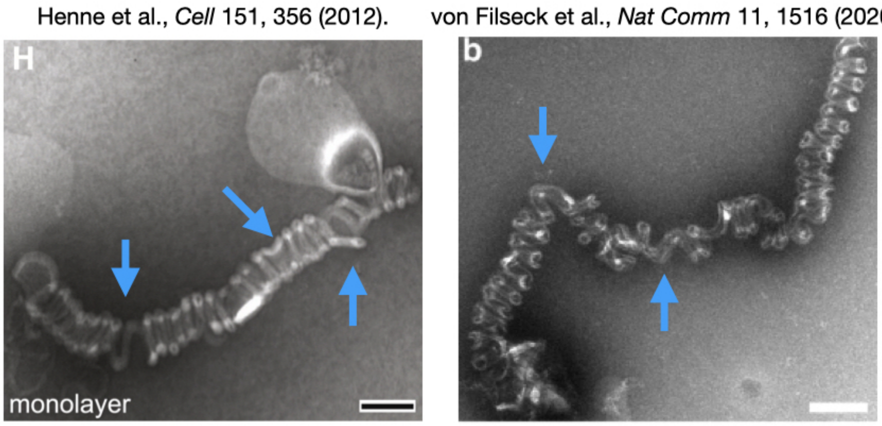


**Figure S7.** Examples of hemihelices reported in eukaryotic (yeast) ESCRT-III filaments. Left panel: Snf7$^{R52E}$/Vps24/Vps2 helices assembled on lipid monolayers, reprinted with permission from Henne et al. [7]. Examples of perversions are marked with blue arrows and scale bar is 100nm. Right panel: Snf7/Vps24/Vps2 filaments grown on large unilamellar vesicles. Reprinted with permission from Von Filseck et al. [8]. Examples of perversions are marked with blue arrows and scale bar is 100nm.

### References

[1] Hongyan Yuan, Changjin Huang, Ju Li, George Lykotrafitis, and Sulin Zhang. One-particle-thick, solvent-free, coarse-grained model for biological and biomimetic fluid membranes. *Phys. Rev. E*, 82:011905, Jul 2010. doi: 10.1103/PhysRevE.82.011905. URL https://link.aps.org/doi/10.1103/PhysRevE.82.011905.

[2] Steve Plimpton, Aidan Thompson, Stan Moore, and Axel Kohlmeyer. Lammps documentation, May 2017. URL http://lammps.sandia.gov/doc/Manual.html.

[3] Alexander Stukowski. Visualization and analysis of atomistic simulation data with ovito — the open visualization tool. *Modelling and Simulation in Materials Science and Engineering*, 18(1):015012, 2010. URL http://stacks.iop.org/0965-0393/18/i=1/a=015012.

[4] Jiangshui Huang, Jia Liu, Benedikt Kroll, Katia Bertoldi, and David R. Clarke. Spontaneous and deterministic three-dimensional curling of pre-strained elastomeric bi-strips. *Soft Matter*, 8:6291–6300, 2012. doi: 10.1039/C2SM25278C.

[5] R.John Ellis. Macromolecular crowding: obvious but underappreciated. *Trends in Biochemical Sciences*, 26(10):597 – 604, 2001. ISSN 0968-0004. doi: https://doi.org/10.1016/S0968-0004(01)01938-7. URL http://www.sciencedirect.com/science/article/pii/S0968000401019387.

[6] F. Zhao. *Confocal Microscopy Tutorial: Lateral And Axial Resolution In Confocal System.*, 2004 (Accessed 6 March 2020). http://www.hi.helsinki.fi/amu/AMU%20Cf_tut/cf_tut_part1-5.htm.

[7] William Mike Henne, Nicholas J. Buchkovich, Yingying Zhao, and Scott D. Emr. The endosomal sorting complex escrt-ii mediates the assembly and architecture of escrt-iii helices. *Cell*, 151(2):356–371, 2018/11/28 2012.

[8] Joachim Moser Von Filseck, Luca Barberi, Nathaniel Talledge, Isabel E Johnson, Adam Frost, Martin Lenz, and Aurélien Roux. Anisotropic escrt-iii architecture governs helical membrane tube formation. *Nature communications*, 11(1):1–9, 2020.
